# Vision is required for the formation of binocular neurons prior to the classical critical period

**DOI:** 10.1101/2021.06.15.448591

**Authors:** Liming Tan, Dario L. Ringach, S. Lawrence Zipursky, Joshua T. Trachtenberg

## Abstract

Depth perception emerges from the development of binocular neurons in primary visual cortex. Vision is required for these neurons to acquire their mature responses to visual stimuli. The prevailing view is that vision does not influence binocular circuitry until the onset of the critical period, about a week after eye opening, and that plasticity of visual responses is triggered by increased inhibition. Here, we show that vision is required to form binocular neurons and to improve binocular tuning and matching from eye opening until critical period closure. Enhancing inhibition does not accelerate this process. Vision soon after eye opening improves the tuning properties of binocular neurons by strengthening and sharpening ipsilateral-eye cortical responses. This progressively changes the population of neurons in the binocular pool and this plasticity is sensitive to interocular differences prior to critical period onset. Thus, vision establishes binocular circuitry and guides binocular plasticity from eye opening.

## Introduction

In mammals with forward facing eyes, the region of visual space in front of the animal is viewed by both eyes. The two eyes are offset in the head, and thus each retina receives slightly different images of visual space. These two visual streams converge on binocular neurons in primary visual cortex which exploit this spatial offset to extract information about the depth of objects in space^1, 2^. To generate a unified perception of the visual world, binocular neurons must integrate inputs from the two eyes with matched receptive field tuning properties^3–5^.

How binocular neurons are established in early postnatal development is not well understood, nor is the role of visual experience in this process. Here, we explore this using 2-photon calcium imaging to track receptive field tuning properties of pyramidal neurons in layer 2/3 in alert mice over the first 4 days after eye opening. We find that vision during this time is required for the establishment of binocular neurons and improves binocular receptive field tuning and matching by changing the population of neurons in the binocular pool much as it does during the classically defined critical period^6^. Moreover, interocular interactions determine the outcome of this early binocular plasticity. Enhancing intracortical inhibition soon after eye opening does not accelerate the emergence of binocular neurons or improve receptive field tuning and matching.

These data revise the conventional view that receptive field tuning of binocular neurons is intrinsically established^7–12^ and that binocular circuitry is unaltered by vision until the later onset of a critical period for binocular plasticity^6, 13–16^. Instead, our measures show that binocular circuitry is informed by vision as soon as the eyes open.

## Results

### Emergence of binocular neurons in the first 4 days after eye opening

To measure the emergence of binocular responses in primary visual cortex we used 2-photon imaging of GCaMP6s responses to measure receptive field tuning of layer 2/3 pyramidal neurons in alert, head-fixed mice over the first 4 days after natural eye opening (**Figure 1A**). In mice, eye opening occurs at or around postnatal day 14 (P14). This period, from P14 to P18, precedes the classically defined opening of the critical period in mice, which begins around postnatal day 21, approximately one week after eye opening^17^. The binocular region of each mouse was identified using retinotopic mapping of GCaMP6s responses, and these maps and the corresponding maps of vasculature were used to target high-resolution 2-photon calcium imaging of single neurons (**Figure 1B**). For each imaged neuron, its receptive field tuning was estimated from the linear regression of the temporally deconvolved calcium response (**Figure 1C**) to a sequence of flashed, high-contrast sinusoidal gratings comprising 18 orientations and 12 spatial frequencies presented at 8 spatial phases for each combination of orientation and spatial frequency^6, 18, 19^ (See also Methods, **Figure S1**). A “tuning kernel” plotting response strength across orientations and spatial frequencies was obtained for each imaged neuron (**Figure 1D**; Figure S1C plots examples of tuning kernels as a function of SNR). Additional examples of tuning kernels are given in **Figure 1E**. Flashed sinusoids were as effective as drifting sinusoids at eliciting cortical responses at P14 (**Figure S2**).

**Figure 1.**
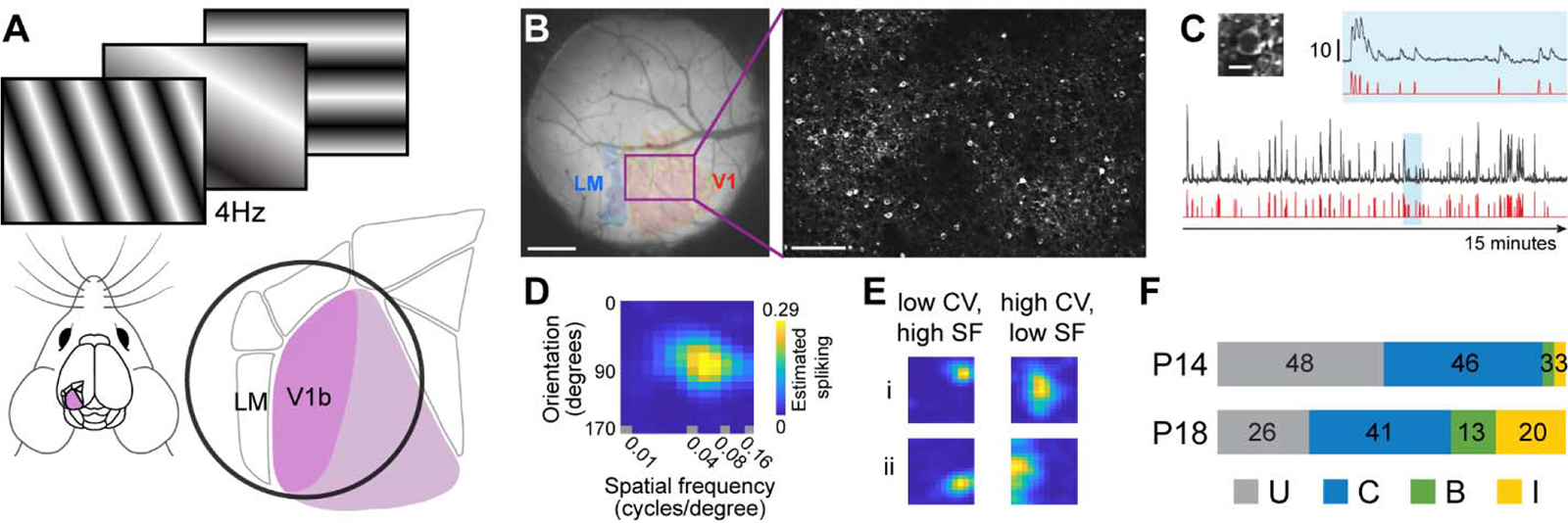
Overview of imaging, receptive field tuning, and development. (A) Schematic overview. Mice passively viewed a series of flashing sinusoidal gratings displayed at 4Hz. Primary visual cortex is highlighted in purple. An expanded diagram of primary visual cortex and higher visual areas highlights the binocular region (V1b) and shows the relative position of the cranial window (black circle). (B) Left: Example image of a cranial window highlighting the binocular visual cortex and the border with LM. The rectangle highlighted in purple delineates the field of view used for of our 2-photon imaging. Scale bar: 0.5 mm. Right: An in vivo example of GCaMP6s labeled pyramidal neurons. Scale bar: 100 μ Image was acquired from a P14 mouse. (C) Left: An image of a single neuron expressing GCaMP6s at P14 (Scale bar: 10 μ. Below). the image is the raw (black) and temporally deconvolved (red) GCaMP6s signal evoked by 15 minutes of visual stimulation. The region in blue is expanded above to show the signal with greater detail. Scale bar: 10 dF/F0. (D) Receptive field tuning kernel of the cell in panel C. Y-axis plots response strength as a function of stimulus orientation. X-axis plots response strength as a function of stimulus spatial frequency, plotted on a log scale. (E) Examples of receptive field tuning kernels from other neurons that show responses that are well tuned to a particular orientation (low circular variance; CV) and relatively high spatial frequency (SF; left column), or that are broadly tuned to a range of orientations (high CV) and relatively lower spatial frequencies (right column). (F) Proportions of imaged cells at P14 (n=2227, 13 mice) and P18 (n=1708, 4 mice) whose receptive field tuning kernels were classified as unresponsive (U), responsive solely to stimulation of the contralateral (C) or ipsilateral (I) eye, or responsive to both eyes (B). See also Figure S1-2.

We found that the rules governing the establishment of binocular neurons after eye opening were similar to those governing this plasticity during the classically defined critical period (P21 to P36)^6^. At eye opening, visually evoked responses were almost entirely driven by contralateral eye stimulation (**Figure 1F**). This low level of innate binocularity is consistent with previous reports^20–23^. Moreover, our measures of contralateral eye evoked receptive field tuning at eye opening are consistent with electrophysiological measures^24^, indicating that GCaMP6s signals at this early age accurately reflect the underlying neurophysiology^25^. Supporting this view, signal-to-noise measures of visually evoked responses at P14 were indistinguishable from those at P16-18 (**Figure S3**). Within the first 4 days after eye opening the fraction of neurons responding solely to stimulation of the ipsilateral eye or to stimulation of either eye (binocular responses) increased 4- to 5-fold, while the fraction of neurons responding to contralateral eye stimulation remained constant (**Figure 1F**).

### Binocular tuning improves via cellular exchange between monocular and binocular pools

To obtain a more informed understanding of the rules governing the emergence of ipsilateral-eye and binocular responses after eye opening, we longitudinally mapped receptive field tuning properties of the same cohort of neurons on P14, P16, and P18 (**Figure 2A**, **Figure S3**-**4**). As during the critical period, the cellular composition of the binocular pool was in continuous flux. Some neurons that were initially monocular gained responsiveness to the other eye and became binocular while other neurons that were initially binocular lost responsiveness to an eye and became monocular (**Figure 2B-E**). **Figure 2B-D** plots examples of neurons with stable receptive field tuning kernels (**Figure 2B**), and those that either gained or lost responsiveness to an eye (**Figure 2C, D**). The trajectories of all 741 longitudinally tracked neurons are plotted in **Figure 2E**. From these trajectories, we find that a majority of binocular neurons form from the conversion of monocular neurons that were initially solely responsive to the contralateral eye (**Figure 2F**).

**Figure 2:**
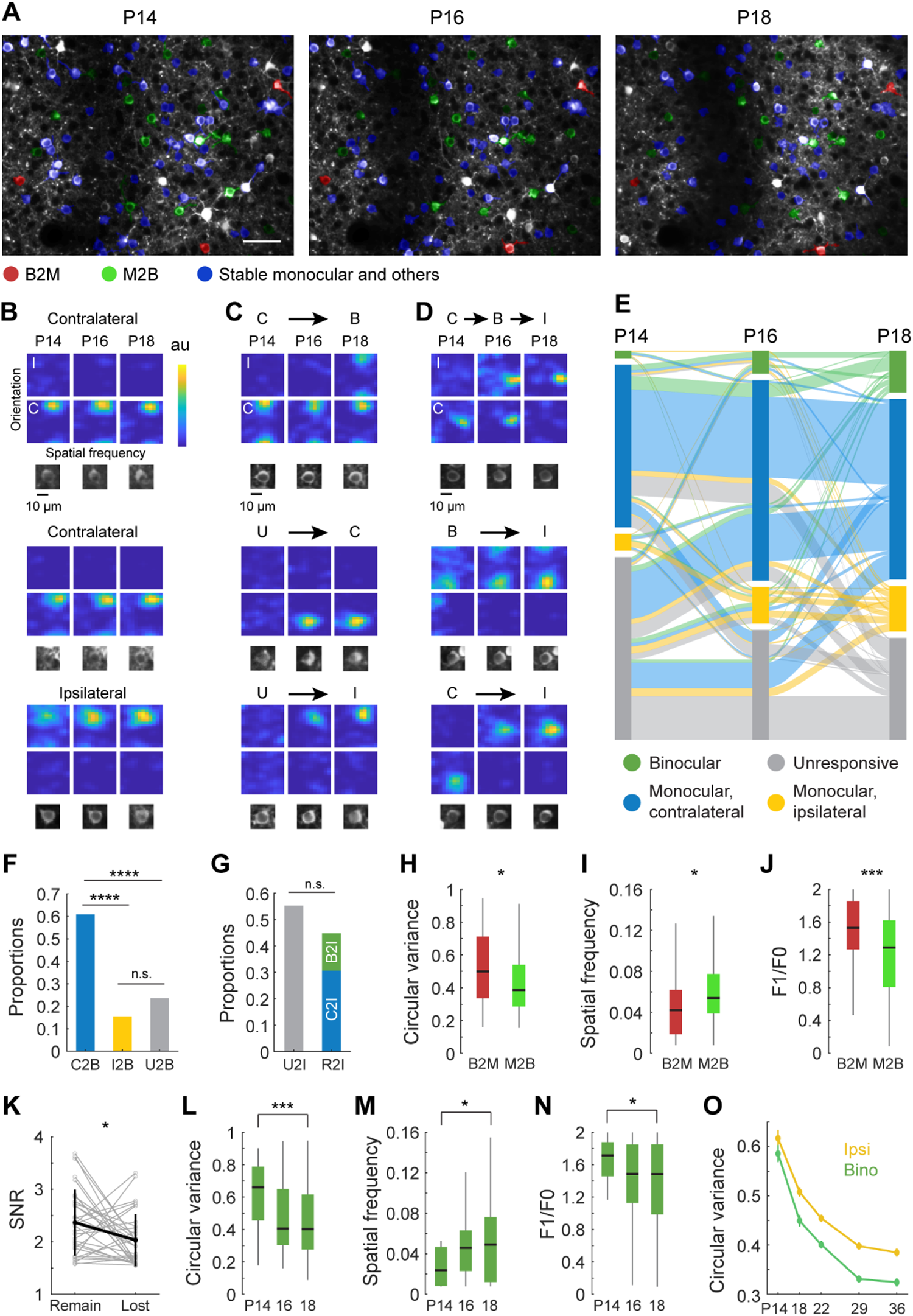
Longitudinal tracking of receptive field tuning properties from eye opening. (A) Example field of view of GCaMP6s expressing pyramidal neurons that was repeatedly imaged on P14, P16, and P18. Neurons with tuning properties tracked across all three time points are colored with red, green and blue masks that represent those that were binocular at P14 and monocular afterwards (red; B2M), monocular at P14 and binocular afterwards (green; M2B), or stable monocular or with other trajectories (blue). Neurons without color masks (white) were not successfully tracked. Scale bar: 50 μm. (B) Examples of receptive field tuning kernels from neurons that were stably monocular, responding solely to either the contralateral (C; upper two rows) or ipsilateral eye (I; bottom row). Below each kernel is an image of the cell. (C) Examples of cells that gained responsiveness to an eye. Top: a cell that was monocularly driven by the contralateral eye at P14 and P16, but gains responsiveness to the ipsilateral eye at P18, thereby becoming binocular (C2B). Middle: a cell that was visually unresponsive at P14 but become solely responsive to contralateral eye stimulation at P16 and remains so at P18 (U2C). Bottom: a cell that was unresponsive at P14 but becomes solely driven by the ipsilateral eye at P16 (U2I). (D) Examples of cells that shift their responsiveness from one eye to the other. Top: a cell that was solely responsive to the contralateral eye at P14, then binocular at P16, then solely responsive to the ipsilateral eye at P18. Middle: a cell that was binocularly responsive at P14, but solely responsive to the ipsilateral eye by P16 and again at P18. Bottom: a cell that was solely responsive to the contralateral eye at P14, but then became solely responsive to the ipsilateral eye thereafter. (E) Sankey diagram plotting the functional trajectory of 741 neurons that were longitudinally imaged at P14, P16, and P18. Length of boxes at each day represents proportions of different groups. Cells are color coded retrospectively. For example, if a neuron ended up binocular at P18, it is color coded green. (F) Proportion of binocular neurons that formed via conversion of neurons in P14-16 or P16-18 that were initially solely responsive to the contralateral eye (C2B, n=67), solely responsive to the ipsilateral eye (I2B, n=17), or unresponsive (U2B, n=26). Chi-square test for pairwise comparisons with Bonferroni correction; ****, p<0.0001 after the Bonferroni correction. (G) Proportion of neurons solely responsive to the ipsilateral eye that formed from initially unresponsive neurons (U2I, n=63), or from neurons that were initially solely driven by the contralateral eye (C2I, n=35) or were binocular (B2I, n=16), and thus responsive to the contralateral eye (R2I). Chi-square test for pairwise comparisons. (H) Circular variance of neurons that were binocular but became monocular (B2M, n=41) and of monocular neurons that became binocular (M2B, n=84). Black horizontal line, median; box, quartiles with whiskers extending to 2.698σ. Mann-Whitney U test; *, p<0.05. (I) As in H, but for measures of spatial frequency preference; *, p<0.05. (J) As in H, but for measures of response linearity; ***, p<0.001. (K) Signal to noise ratio (SNR) of responses to each eye in binocular neurons prior to the loss of one eye’s responses (n=33). Gray, measurements for individual neurons; black, mean ± standard deviation. Mann-Whitney U test of SNR between remained and lost eye specific responses; *, p<0.05. Note: the SNR of the response of the eye that remained was stronger than that of the eye that was lost. (L) Circular variance of binocular neurons measured at P14 (n=15), P16 (n=46), and P18 (n=84). Mann-Whitney U test between P14 and 18; ***, p<0.001. Note the gradual improvement in orientation selectivity with age. (M) As in L, but for spatial frequency tuning; *, p<0.05. (N) As in L, but for linearity. *, p<0.05. (O) Plot of circular variance for binocular neurons (n=74, 220, 450, 417, 339) and neurons driven by the ipsilateral eye (n=147, 562, 1024, 931, 832) from eye opening until the end of the critical period at P36. Note the continual improvement with age. See also Figure S3-5.

The emergence of monocular, ipsilateral-eye cortical responses occurred via three routes. In order of prevalence, one was via conversion of previously unresponsive neurons (**Figure 2G**, U2I, 55%; see also **Figure 2**), a second was via conversion of neurons that were initially solely responsive to the contralateral eye (C2I, 30%), and the third was via conversion of binocular neurons (B2I, 15%; **Figure 2G**; see also **Figure 2D, E**).

As during the critical period^6^, the exchange of neurons to and from the binocular pool in the first four days after eye opening was correlated with receptive field tuning. Monocular neurons that became binocular had more selective orientation tuning, preferred higher spatial frequency stimuli, and were more complex than binocular neurons that became monocular (**figure 2H-J**). When binocular neurons lost responsiveness to one eye, it was the weaker eye that was eliminated (**figure 2K**). Moreover, cells that switched their preference from one eye to the other (e.g. C2I or I2C) were not as sharply tuned as those that became binocular (C2B or I2B: CV=0.45 ± 0.22, n=84; C2I or I2C: CV=0.55 ± 0.19, n=53; mean ±std; P=0.0010).

This gain and loss of binocular neurons as a function of receptive field tuning from P14 to P16 to P18 resulted in a progressive improvement of binocular receptive field tuning (**Figure 2L-N**). This was confirmed by comparing acute measures of receptive field tuning of 74 binocular neurons at P14 (of 2227 imaged neurons) and 220 at P18 (of 1708 imaged neurons) (**Figure S5**). Taken together, these measures show that the rules governing the establishment of binocular neurons and improvements in their tuning in the first four days after eye opening are similar to those during the classically defined critical period^6^. Thus, from eye opening to P36, there is a continual improvement in binocular tuning that is paralleled by improvements in the tuning of cortical responses to ipsilateral-eye stimulation. This parallel improvement is plotted in **Figure 2O**, which includes data from P22 to P36 that was previously reported in Tan et al., 2020.

### Vision is required to establish binocular responses immediately after eye opening

To determine the role of early vision in the emergence of ipsilateral-eye and binocular responses, we reared mice in darkness from P12 to P18 and then measured receptive field tuning properties evoked via stimulation of either eye. In dark reared mice, about half the normal number of binocular neurons formed and 2/3^rd^ of the normal number of neurons were responsive to stimulation of the ipsilateral eye (**figure 3A-C**). The fraction of neurons responsive to stimulation of the contralateral eye, however, was unaffected (**Figure 3A, D**). The few binocular neurons that formed in these dark reared mice had much poorer receptive field tuning properties and binocular matching coefficients than normal (**Figure 3E-I**). This reflected the poor tuning of ipsilateral-eye responses in these dark reared mice (**Figure 3J,K**). Thus, early vision is required for the normal emergence and sharpening of cortical responses to ipsilateral-eye stimulation^26, 27^ and this drives similar increases in the fraction of binocular neurons and improvements in their receptive field tuning properties. Again, this is similar to what occurs during the classically defined critical period when vision improves binocular tuning via its refinement of the ipsilateral-eye pathway.

**Figure 3.**
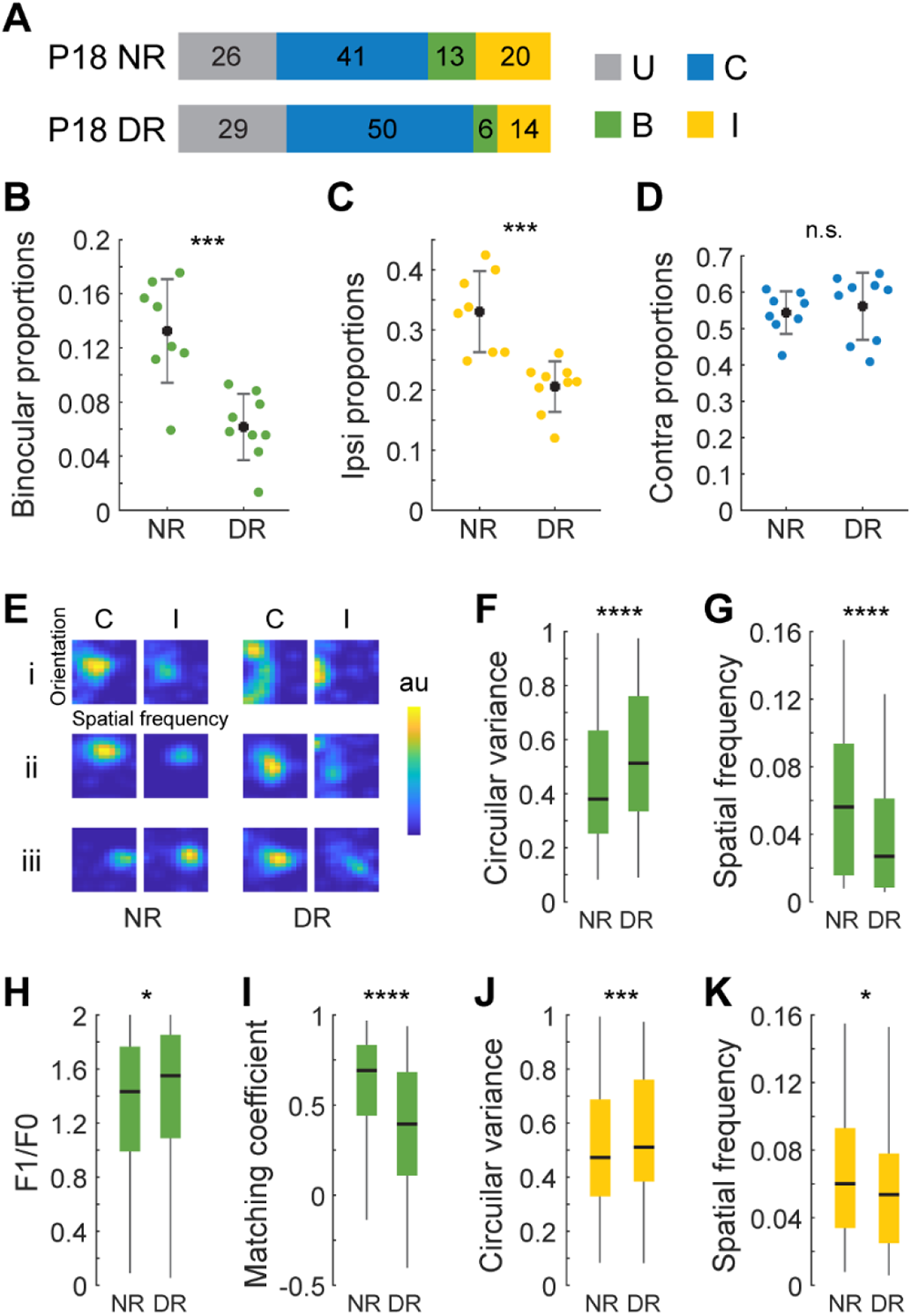
Vision is required for the emergence and sharpening of cortical responses to ipsilateral eye stimulation. (A) Proportion of neurons in P18 mice that are solely responsive to the ipsilateral (I) or contralateral (C) eye or are binocular (B) in normally reared (NR, n=4) and dark reared (DR, n=4) mice. Color coding is defined in the adjacent legend. U represents unresponsive neurons. (B) Fraction of binocular neurons as a function of rearing condition. Each point is from a single imaging plane. Mean and standard deviation are shown as black dots and lines. Mann-Whitney U test; ***, p<0.001. (C) As in B but for neurons responding to ipsilateral eye stimulation; ***, p<0.001. (D) As in B but for neurons responding to contralateral eye stimulation. Note the absence of an effect of dark rearing. (E) Examples of receptive field tuning kernels for binocular neurons in normally reared and dark reared mice. Examples of kernels evoked via contralateral or ipsilateral eye stimulation for the same neuron are plotted adjacent to each other. Kernels from 3 example neurons representing the 1st, 2nd, and 3rd quartiles in the distribution of tuning quality (joint measurements of circular variance, spatial frequency preference, complexity and binocular matching coefficient) in each condition are labeled i-iii. (F) Boxplots of the distribution of circular variance values recorded from binocular neurons in normally reared (n=220 cells, 4 mice) and dark reared (n=115 cells, 4 mice) mice. Black horizontal line, median; box, quartiles with whiskers extending to 2.698σ Mann-Whitney U test; ****, p<0.0001. (G) As in F but for measures of spatial frequency tuning; ****, p<0.0001. (H) As in F but for measures of complexity; *, p<0.05. (I) As in F but for measures of binocular matching coefficients; ****, p<0.0001. Note the poorer matching in the absence of vision. (J) Boxplots of circular variance values recorded in response to stimulation of the ipsilateral eye in normally reared (n=562 cells, 4 mice) and dark reared (n=367 cells, 4 mice) mice. Black horizontal line, median; box, quartiles with whiskers extending to 2.698σ. Mann-Whitney U test; ***, p<0.001. (K) Same as J but for measures of spatial frequency tuning; *, p<0.05. See also Figure S6.

### Binocular plasticity begins at eye opening

Our longitudinal imaging data indicate that the emergence of cortical responses to ipsilateral-eye stimulation is governed, in part, by binocular interactions where neurons that were initially responsive to the contralateral eye become solely responsive to the ipsilateral eye (cf. **Figure 2D, G**). We examined this more directly using the classic paradigm of monocular deprivation. Binocular plasticity in primary visual cortex is studied by comparing the effects of monocular deprivation, which creates a strong imbalance in the ability of one eye to drive cortical responses, to those of binocular visual deprivation where both eyes are equally impaired in their ability to drive cortical responses. Here, we compare the effects of monocular lid suture to those of dark rearing. Dark rearing reveals the impact of vision on the development of visually evoked responses^28^, while monocular deprivation reveals the differential impact of interocular interactions^26^. We avoid the use of binocular lid suture because it might introduce interocular differences in activity. Light transmission through the two sutured eyelids can differ by as much as 1 log unit^29^.

The effects of monocular deprivation were distinct from those of dark rearing. We found that suturing the ipsilateral eye from P14 to P18 was far more detrimental to the development of ipsilateral-eye responses than was dark rearing from P14 to P18 (**Figure 4A-E**). By comparison to normally reared mice, unilateral deprivation of the ipsilateral eye resulted in both a reduction in the pool of neurons responsive to ipsilateral-eye stimulation (**Figure 4A, B**; 70% reduction) and an expansion of the contralateral pool (**Figure 4A, F**; 20% expansion). Dark rearing, however, was less deleterious; the ipsilateral pool was reduced by 46% (**Figure 4A, B**) and the size of the contralateral pool was unaffected (see **Figure 3D**). Similarly, the normal improvement of receptive field tuning of ipsilateral-eye responses was completely suppressed by unilateral deprivation of the ipsilateral eye, but less affected by dark rearing (**Figure 4C-E**). By contrast, the development of receptive field tuning of contralateral eye responses was not impaired by deprivation of the ipsilateral eye (**Figure 4G-J**). Thus, the normal emergence and refinement of ipsilateral-eye tuning after eye opening is both vision dependent and highly sensitive to interocular differences in visual experience.

**Figure 4:**
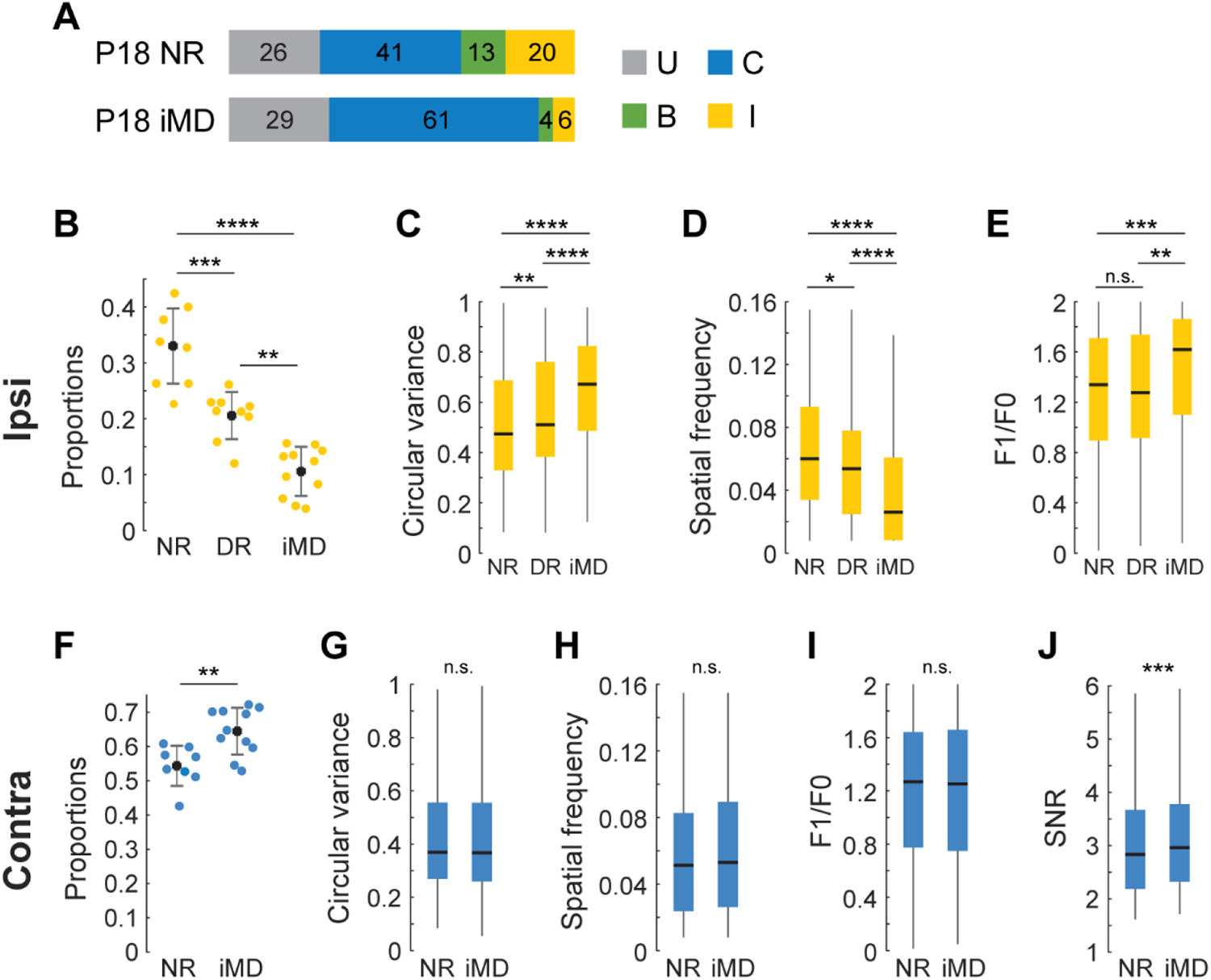
The development of ipsilateral eye cortical responses requires vision and is sensitive to monocular deprivation. (A) As in Figure 3A, but for normally reared (NR, n=4) and ipsilateral eye sutured (iMD, n=4) mice. (B) Plot of the proportion of cells that respond to visual stimulation of the ipsilateral eye in P18 mice that were normally reared (NR, n=4), dark reared (DR, n=4), or that experienced monocular lid suture of the ipsilateral eye (iMD, n=4) from P14 to P18. Each point is from a single imaging plane. Mean and standard deviation are shown as black dots and lines. Mann-Whitney U test with Bonferroni correction. **, p<0.01; ***, p<0.001; ****, p<0.0001 after correction. (C-E) Boxplots of measures of circular variance, spatial frequency preference, and complexity of neuron responses evoked by ipsilateral eye stimulation in NR (n=562 cells), DR (n=367 cells), and iMD (n=178 cells) mice. Black horizontal line, median; box, quartiles with whiskers extending to 2.698σ. Mann-Whitney U test with Bonferroni correction. *, p<0.05; **, p<0.01; ***, p<0.001; ****, p<0.σ0001 after correction. (F-J) Same as in B-E, but for responses of neurons evoked via contralateral eye stimulation in NR (n=916 cells) and iMD (n=1104) mice. Note the quality of contralateral eye tuning is not affected by iMD. See also Figure S6.

Interocular interactions did not influence the development of cortical responses to contralateral eye stimulation (**Figure 5A-E**). The proportion of neurons responding to contralateral eye stimulation and the receptive field tuning measures of these responses were indistinguishable between mice exposed to unilateral lid suture of the contralateral eye from P14 to P18 and those kept in the dark from P14 to P18 (**Figure 5A-E**). Paradoxically, deprivation of the contralateral eye at these ages impaired the maturation of cortical responses to ipsilateral-eye stimulation (**Figure 5F-J**). Although the fraction of neurons responding to ipsilateral-eye stimulation and the sharpness of their orientation tuning remained similar (**Figure 5F, G**), the spatial frequency preferences of their responses, their complexity and signal to noise ratio were 33%, 15% and 10% lower than normal, respectively (**Figure 5H-J**). These measures provide additional evidence that interocular differences in visual experience in the first few days after eye opening influence the development of cortical responses to the ipsilateral eye, and thus binocular responses.

**Figure 5:**
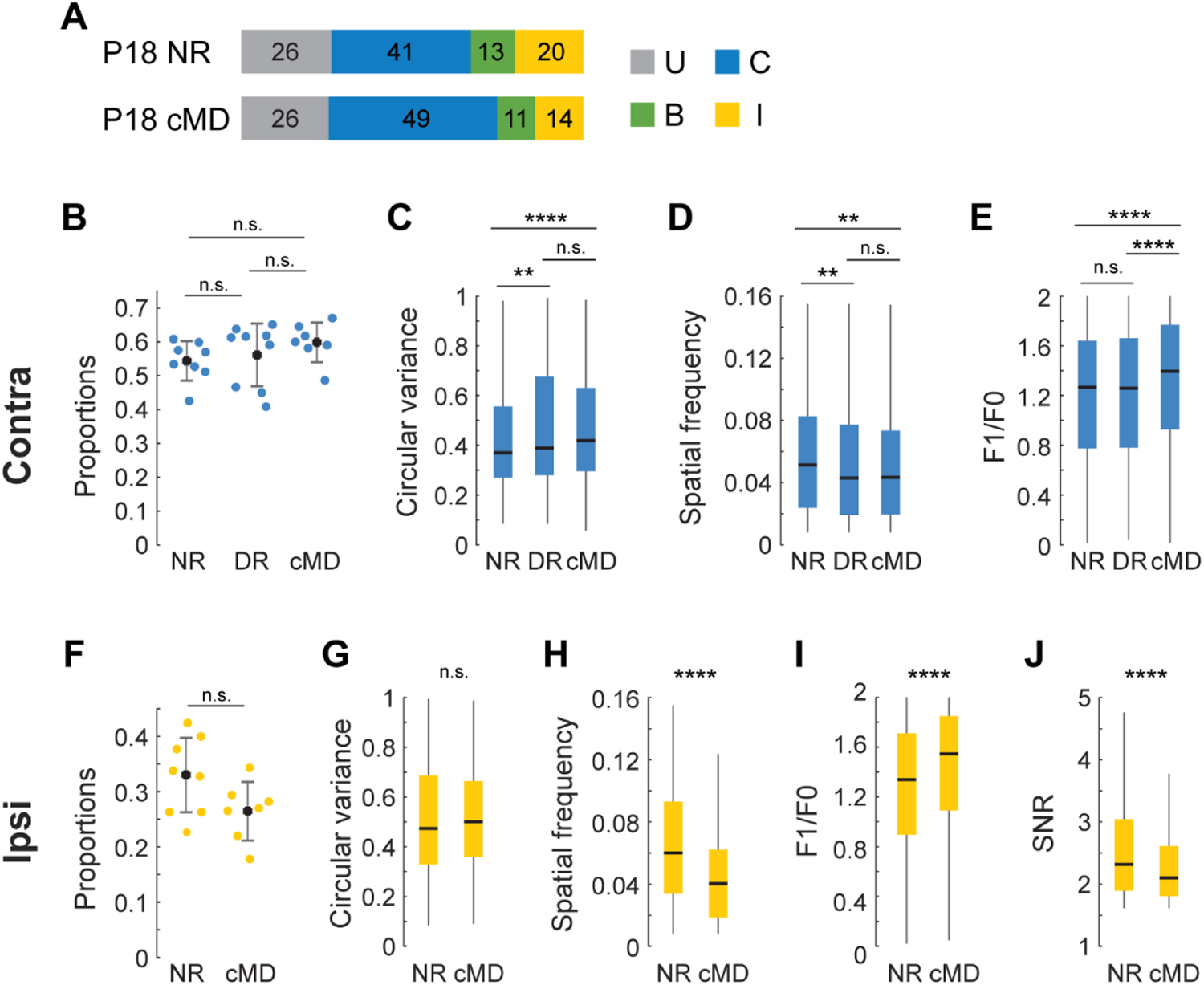
The development of ipsilateral eye cortical responses from P14 to P18 is affected by contralateral eye visual experience. (A) As in Figure 4A, but for normally reared (NR, n=4) and contralateral eye sutured (cMD, n=3) mice. (B) Plot of the proportion of cells that respond to visual stimulation of the contralateral eye in P18 mice that were normally reared (NR, n=4), dark reared (DR, n=4), or that experienced monocular lid suture of the contralateral eye (cMD, n=3) from P14 to P18. Each point is from a single imaging plane. Mean and standard deviation are shown as black dots and lines. Mann-Whitney U test with Bonferroni correction. (C-E) Boxplots of measures of circular variance, spatial frequency preference, and complexity of neural responses evoked by contralateral eye stimulation in NR (n=916 cells), DR (n=1004 cells), and cMD (n=1215 cells) mice. Black horizontal line, median; box, quartiles with whiskers extending to 2.698 σ . Mann-Whitney U test with Bonferroni correction. **, p<0.01; ****, p<0.0001 after correction. (F-J) Same as in B-E, but for responses of neurons evoked via ipsilateral eye stimulation in NR (n=562) and cMD (n=507) mice. Note the decrease in ipsilateral eye tuning quality in cMD mice. See also Figure S6.

### Enhancing inhibition does not accelerate the development of binocular tuning

Fast-spiking inhibition mediated by parvalbumin-expressing interneurons is not measurable within the first 3 days after natural eye opening^24^. Maturation of this inhibition triggers the opening of the classically defined critical period for monocular deprivation and is thought to be needed for visual experience to shape binocular circuitry^16, 30^. Pharmacological enhancement of fast-spiking inhibition immediately after eye opening, either via administration of the use-dependent GABA-A agonist Diazepam^14, 15, 31, 32^, or via over-expression of neurotrophic factors that drive the development of parvalbumin-expressing interneurons results in a precocious opening of the critical period^33, 34^.

To test whether precocious development of fast-spiking inhibition accelerates the influence of vision on binocular circuitry, we measured receptive field tuning evoked via stimulation of each eye in P18 mice that received intraperitoneal injections of Diazepam on P15 and P16 (30 mg kg^-1^)^35–37^. We found that enhancing inhibition impeded rather than accelerated binocular development. Binocular neurons formed in normal numbers (**Figure 6A**), but receptive field tuning was impaired (**Figure 6B-E**). Although responses evoked via stimulation of the contralateral eye in these binocular neurons were normal, ipsilateral-eye evoked tuning was characterized by lower spatial frequency preferences and simpler responses (**Figure 6B, D, E**). This differential impact on ipsilateral-eye tuning resulted in the establishment of binocular neurons with lower binocular matching coefficients (**Figure 6B, C**). Thus, accelerating the onset of cortical inhibition does not accelerate the normal maturation of binocular neurons with complex and matched receptive field tuning^37^.

**Figure 6.**
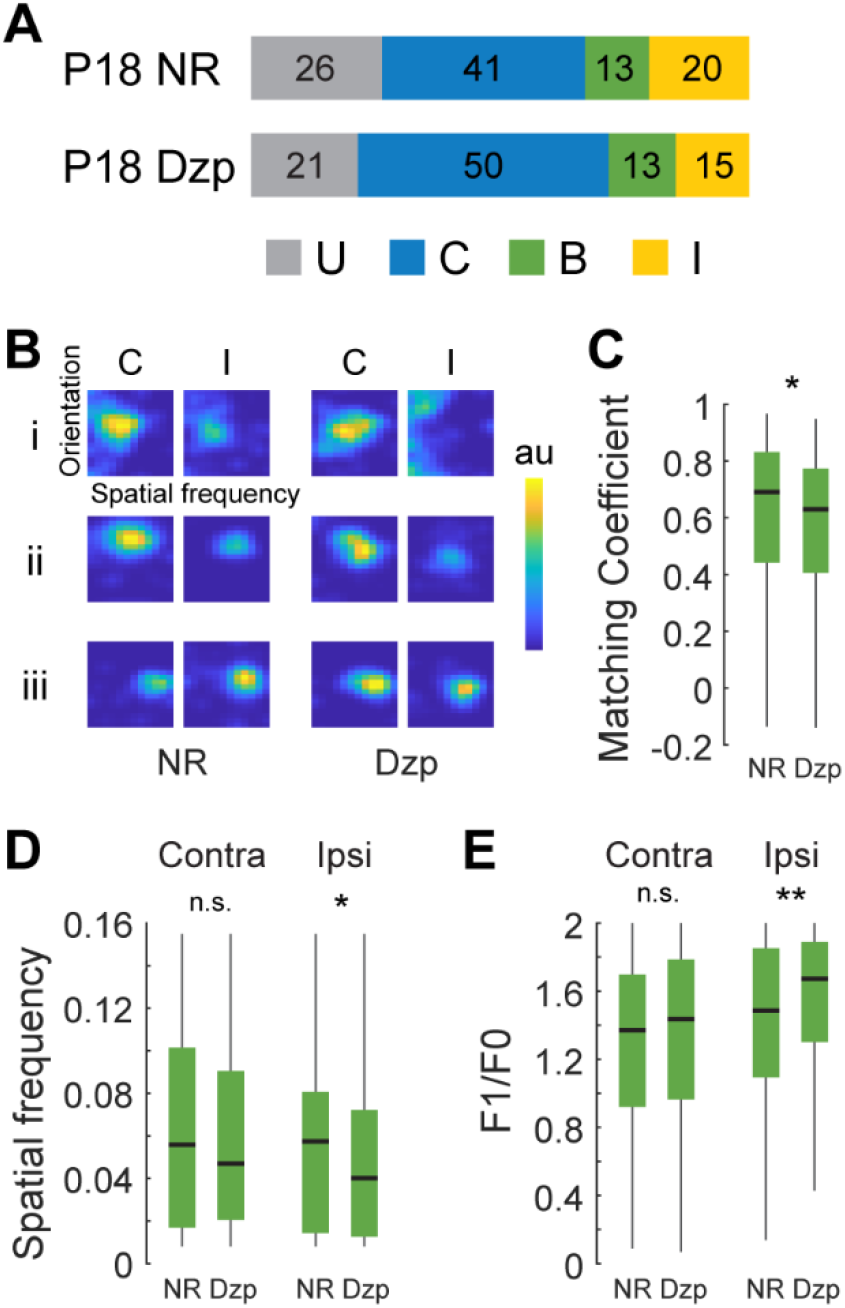
Enhancing early inhibition does not accelerate the development of binocular neurons. (A) Proportions of imaged cells at P18 whose receptive field tuning kernels were classified as unresponsive (U), responsive solely to stimulation of the contralateral (C) or ipsilateral (I) eye, or responsive to both eyes (B) in normally reared mice (NR, n=4 mice) and in mice that received I.P. injections of diazepam (Dzp, n=4 mice) on P15 and P16. (B) Examples of receptive field tuning kernels for binocular neurons in normally reared and diazepam treated mice. Examples of kernels evoked via contralateral or ipsilateral eye stimulation for the same neuron are plotted adjacent to each other. Kernels from 3 example neurons representing the 1st, 2nd, and 3rd quartiles of the tuning quality distribution for each condition are labeled i-iii, as in Figure 3E. (C) Boxplot of binocular matching coefficients in NR mice (n=4 mice, 220 cells) and Dzp treated mice (n=4 mice, 277 cells). Black horizontal line, median; box, quartiles with whiskers extending to 2.698σ. Mann-Whitney U test, *, p<0.05. (D) Boxplots of spatial frequency preferences of binocular neurons evoked via contralateral or ipsilateral eye stimulation in NR and Dzp mice. Black horizontal line, median; box, quartiles with whiskers extending to 2.698σ. Mann-Whitney U test, *, p<0.05 for Ipsi. (E) As in D, but for complexity, **, p<0.01 for Ipsi. See also Figure S6.

## Discussion

The opening of the critical period has been defined by measuring the effects of brief periods of monocular deprivation applied at different times after eye opening on cortical ocular dominance^11, 17, 38^. These brief periods of monocular deprivation result in the loss of cortical responsiveness to vision through the deprived eye, thus shifting ocular dominance measures of binocular responses toward the open eye. In carnivores and rodents, the onset of the critical period for the effects of unilateral lid suture on ocular dominance begins somewhat suddenly about a week after eye opening. This is triggered by the maturation of intracortical inhibition mediated by fast-spiking, parvalbumin-expressing interneurons^9, 30, 39, 40^. Supporting this, the onset of the critical period can be temporally shifted by manipulating this inhibitory maturation^14, 15, 33, 34^. Emergent from these measures is the view that vision-dependent cortical plasticity is not engaged until fast-spiking inhibition has matured. Thus, vision drives cortical responses at eye opening but cannot shape binocular circuitry until about a week later.

An extension of this view is that binocular circuitry is intrinsically established by molecular guidance cues and patterns of spontaneous activity^9^. Supporting this, anatomical measures of the periodicity of ocular dominance columns in carnivores and primates (mice do not have ocular dominance columns) show that this spacing is adult-like at eye opening^12, 41^ and is established even in the absence of retinal input^42^ (though, in this study, On/Off clustering may have been misconstrued for ocular dominance). These measures give rise to the notion that vision is necessary for the maintenance but not the establishment of binocularity.

By contrast, we demonstrate here that early vision is required for the formation of normal numbers of binocular neurons and to improve binocular cell receptive field tuning and matching. In agreement with previous studies, our measures show that few binocular neurons are present at eye opening^20–22, 41^. Within the first four days thereafter, a near-adult number of binocular neurons is established. This occurs through the vision-dependent strengthening and refinement of cortical responses to the ipsilateral eye. In agreement with earlier data obtained using intrinsic signal optical imaging, we find that the development of ipsilateral-eye responses is sensitive to vision through the ipsilateral eye, to vision through the contralateral eye, and to interocular differences therein^27^. Additionally, the continuous improvement of ipsilateral-eye cortical responses drives the early exchange of neurons between monocular and binocular pools as a function of receptive field tuning and binocular matching much as it does during the classically defined critical period^6^.

One caveat of dark-rearing experiments is that dark rearing delays the normal progression of synaptic and physiological changes that are triggered by eye opening^43–45^. While we did not observe any decrement in signal to noise measures of contralateral-eye evoked responses with dark rearing, any changes in circuit excitability due to dark rearing must be considered.

The maturation of fast-spiking inhibition is thought to trigger the onset of the critical period for binocular plasticity in primary visual cortex. Consistent with this view, visually-evoked responses of fast-spiking interneurons have not been recorded within the first 4 days after natural eye opening in mice^24^ – prior to the onset of the critical period. However, binocular plasticity is very much engaged during this pre-critical period. Binocular matching proceeds via vision-dependent plasticity, and brief monocular deprivation of the ipsilateral eye drives the loss of ipsilateral eye responses and the expansion of contralateral eye responses. Thus, vision-dependent binocular plasticity does not require the engagement of fast-spiking inhibition. Indeed, pharmacological enhancement of inhibition during this pre-critical period did not accelerate any aspect of the normal development of binocular responses. Instead, it only seemed to impair the development of ipsilateral and binocular responses. A potential concern is that diazepam administration is followed by a period of severe drowsiness. This may impact changes in binocular tuning that take place on a very fast time scale. This view that increased inhibition is detrimental to the development of cortical responses to the ipsilateral eye is consistent with conclusions drawn from recent work showing that an excess number of parvalbumin-expressing interneurons in cortex, and thus excess inhibition, impairs the normal development of ipsilateral-eye responses and binocular vision^46^. Thus, while other forms of early inhibition may shape receptive fields of binocular neurons at a more fine-scaled level, fast-spiking inhibition does not appear to be needed for vision to influence the establishment of binocular neurons, the process of binocular matching, or shifts in cortical binocularity following monocular deprivation of the ipsilateral eye, as all of these occur during the pre-critical period.

By contrast to what we found for the ipsilateral eye, the development of cortical responses to the contralateral eye matures fairly well even in the absence of vision. Thus, cortical responses to contralateral eye stimulation are, largely, intrinsically established^47, 48^. Cortical responses to the contralateral eye gain a requirement for vision about a week after eye opening, when, for the first time, deprivation of the contralateral eye results in the loss of cortical responsiveness to that eye^17^. A large body of work shows that this is triggered by the maturation of fast-spiking inhibition^9, 10, 14, 33, 34^.

Remarkably, all studies of early ocular dominance plasticity in mice prior to P21 appear to have used lid suture of the contralateral eye. Our measures show that cortical responses to this eye are intrinsically established and insensitive to vision prior to at least P18. Had previous monocular deprivation experiments examined the effects of ipsilateral-eye deprivation, the requirement of early vision for the emergence of ipsilateral-eye cortical responses and the improvement in binocular receptive field tuning would have been revealed.

The data we present revise our understanding of the critical period for binocular plasticity in primary visual cortex. We show that binocular circuitry is not intrinsically established but requires vision during the first several days after eye opening. Moreover, visual experience-dependent cortical plasticity does not begin at the onset of the critical period, it begins at eye opening. The opening of the classically defined critical period, therefore, does not usher in a period when visual cortical circuitry is suddenly susceptible to visual experience – this susceptibility begins immediately. Instead, the onset of the classical critical period appears to usher in a period when intrinsically established contralateral eye evoked cortical responses require vision for their maintenance; this is triggered by the maturation of fast-spiking inhibition. These data shift the position of fast-spiking inhibition from one in which it is centrally involved in transforming visual experience into circuit changes to one in which it is involved in maintaining and consolidating contralateral eye cortical responses. The idea that intracortical inhibition consolidates circuitry and that suppressing inhibition promotes cortical plasticity is well established^49–51^. Taken together, our data provide a new framework for the role of early visual experience in the development of binocular circuitry.

## Acknowledgements

We thank all members of the Zipursky lab and Trachtenberg lab for constructive criticism of the manuscript. We thank Asavari Tiku for mouse genotyping. We thank UCLA DLAM for their assistance with diazepam injections. We thank Lila Trachtenberg for providing the drawing in Figure 1A. This study was funded by NIH R01EY023871 and NIH R01 EY027407 (J.T.T.) and NIH R01 NS116471 and NIH EB022915 (D.L.R.). We thank W. M. Keck Foundation for funding this project. S.L.Z. is an investigator of Howard Hughes Medical Institute.

## Author Contributions

Conceptualization, L.T., J.T.T., S.L.Z. and D.L.R.; Methodology, L.T., J.T.T., and D.L.R.; Investigation and Analysis, L.T.; Writing – Original Draft, L.T. and J.T.T.; Writing – Review & Editing, L.T., J.T.T., S.L.Z. and D.L.R.; Funding Acquisition, J.T.T., S.L.Z. and D.L.R.; Resources, J.T.T., S.L.Z.; Supervision, J.T.T.

---

**Figure S1.**
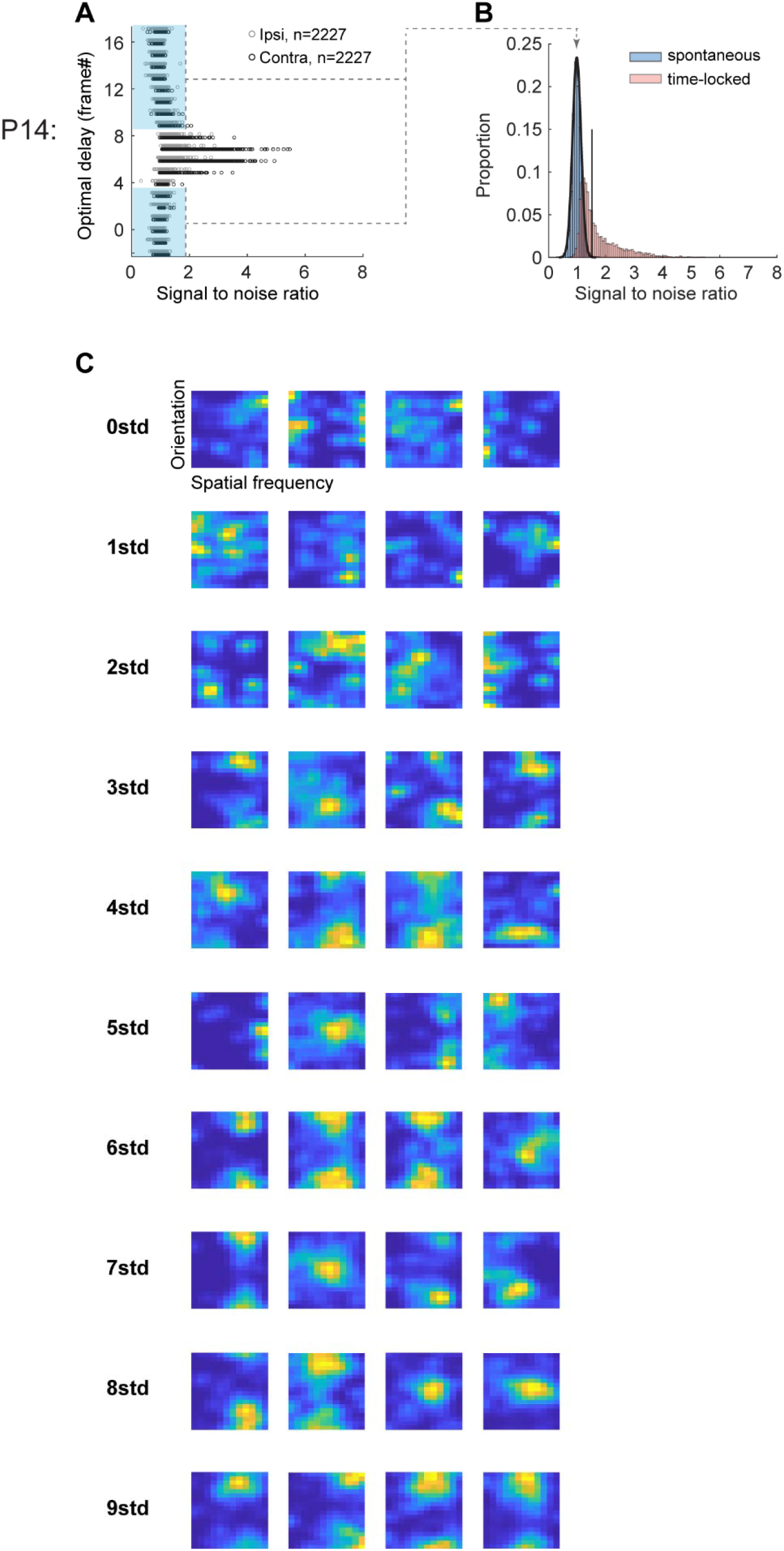
SNR distribution of L2/3 neurons acutely imaged from 13 mice at P14, and examples of receptive field tuning kernels as a function of SNR. Related to Figure 1. (A) Plot of SNR as a function of optimal delay for all 2227 L2/3 neurons imaged in P14 mice. Dark and light gray represent responses to contralateral and ipsilateral eye stimulation, respectively. A cell with an optimal delay of -2 to 3 or 9 to 17 frames post stimulus onset was considered to be visually unresponsive (blue shading). (B) Blue shaded histogram and normal distribution fit of SNR values for unresponsive neurons. Black vertical line is 3 standard deviations above the mean of this normal distribution. Red shaded histogram: neurons whose spiking occurred between 4 and 8 frames post stimulus. Visually responsive neurons are those to the right of the vertical line, with optimal delays in this window that also had an SNR value above the threshold. (C) Examples of tuning kernels as a function of SNR. Each row plots tuning kernels of four example neurons, whose SNR values are 0,1,2…9 standard deviations above noise mean. Significant tuning kernels should appear close to Gaussian distribution along the X- and Y-axis. SNR values below 3 standard deviations of noise mean are unstructured.

**Figure S2.**
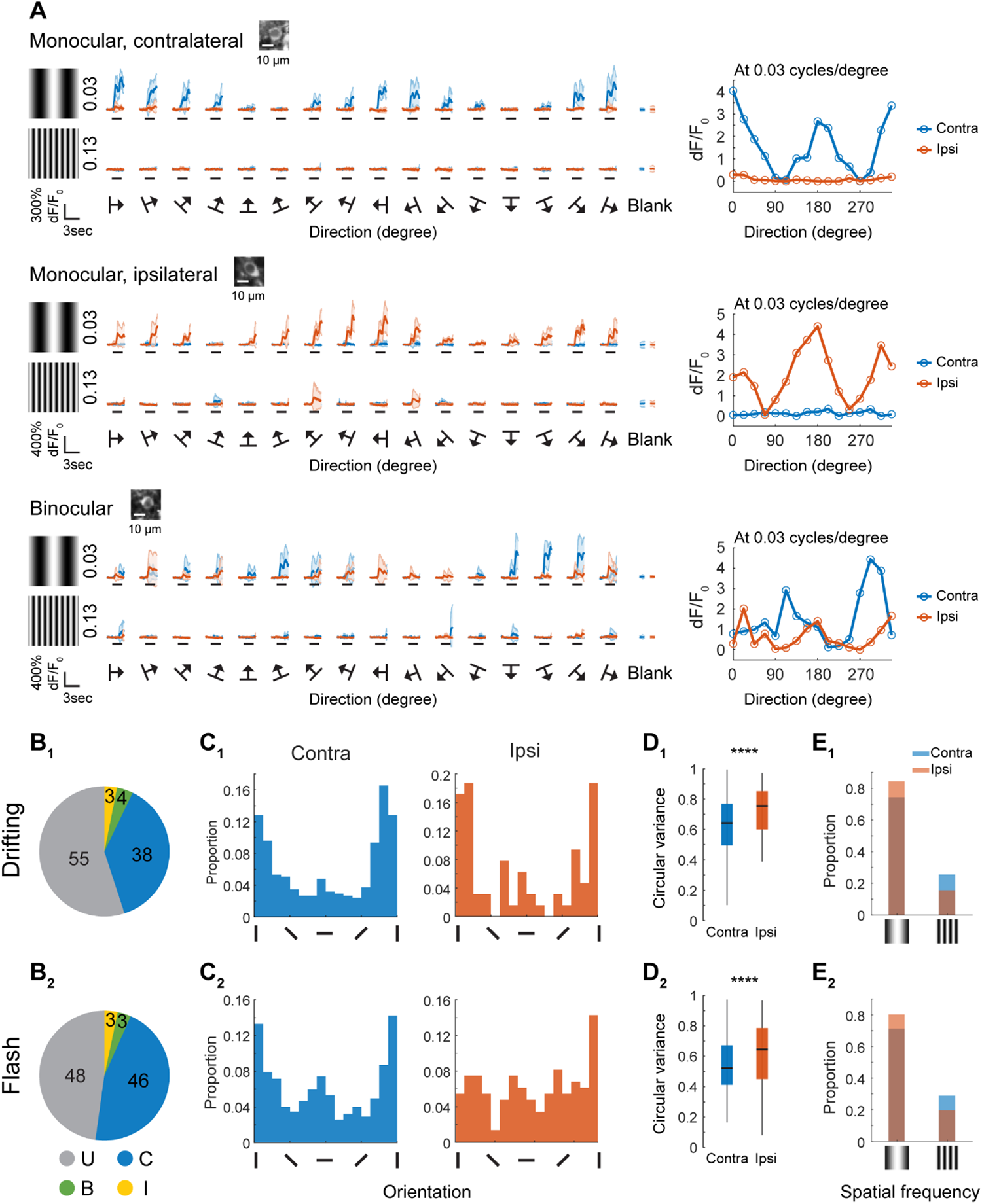
Tuning measurements using drifting grating and flash grating were in close agreement at P14. Related to Figure 1. (A) Response traces (left panel, mean ± standard deviation of 5 trials for each stimulus) and orientation tuning curves to trial-averaged responses at optimal spatial frequency (right panel, mean) for a monocular contralateral, monocular ipsilateral and binocular neuron in P14 mice. The image of the soma for each neuron is shown on top of each left panel. The black line under each trace indicates visual stimulation (2 sec). Stimuli were drifting gratings at 16 evenly spaced directions at 2 spatial frequencies (0.03 and 0.13 cycles/degree). Note for all three cells, response signals at preferred stimuli showed modulation at the frequency of drifting gratings (1 Hz). (B) Proportions of imaged pyramidal neurons in layer 2/3 at P14 using drifting (B1, n=896, 3 mice) or flashed gratings (B2, n=2227, 13 mice). Neurons were classified as unresponsive (U), responsive solely to stimulation of the contralateral (C) or ipsilateral (I) eye, or responsive to both eyes (B). (C) Histograms of orientation preference evoked via contralateral eye or ipsilateral eye stimulation using drifting (C1, Contra, n=375; Ipsi, n=64) or flash gratings (C2, Contra, n=1089; Ipsi, n=147). (D) Boxplots of circular variance to the contralateral eye or ipsilateral eye stimulation using drifting (D1, Contra, n=375; Ipsi, n=64) or flashed gratings (D2, Contra, n=1089; Ipsi, n=147). Black horizontal line, median; box, quartiles with whiskers extending to 2.698 σ. Mann-Whitney U test, ****, p<0.0001. (E) Bar graphs showing proportions of neurons preferring low (0.03 cycles/degree) or high spatial frequencies (0.13 cycles/degree) using drifting (E1, Contra, n=375; Ipsi, n=64) or flashed gratings (E2, Contra, n=1089; Ipsi, n=147). The cutoff between low and high spatial frequencies for flash grating data is 0.65 cycles/degree.

**Figure S3.**
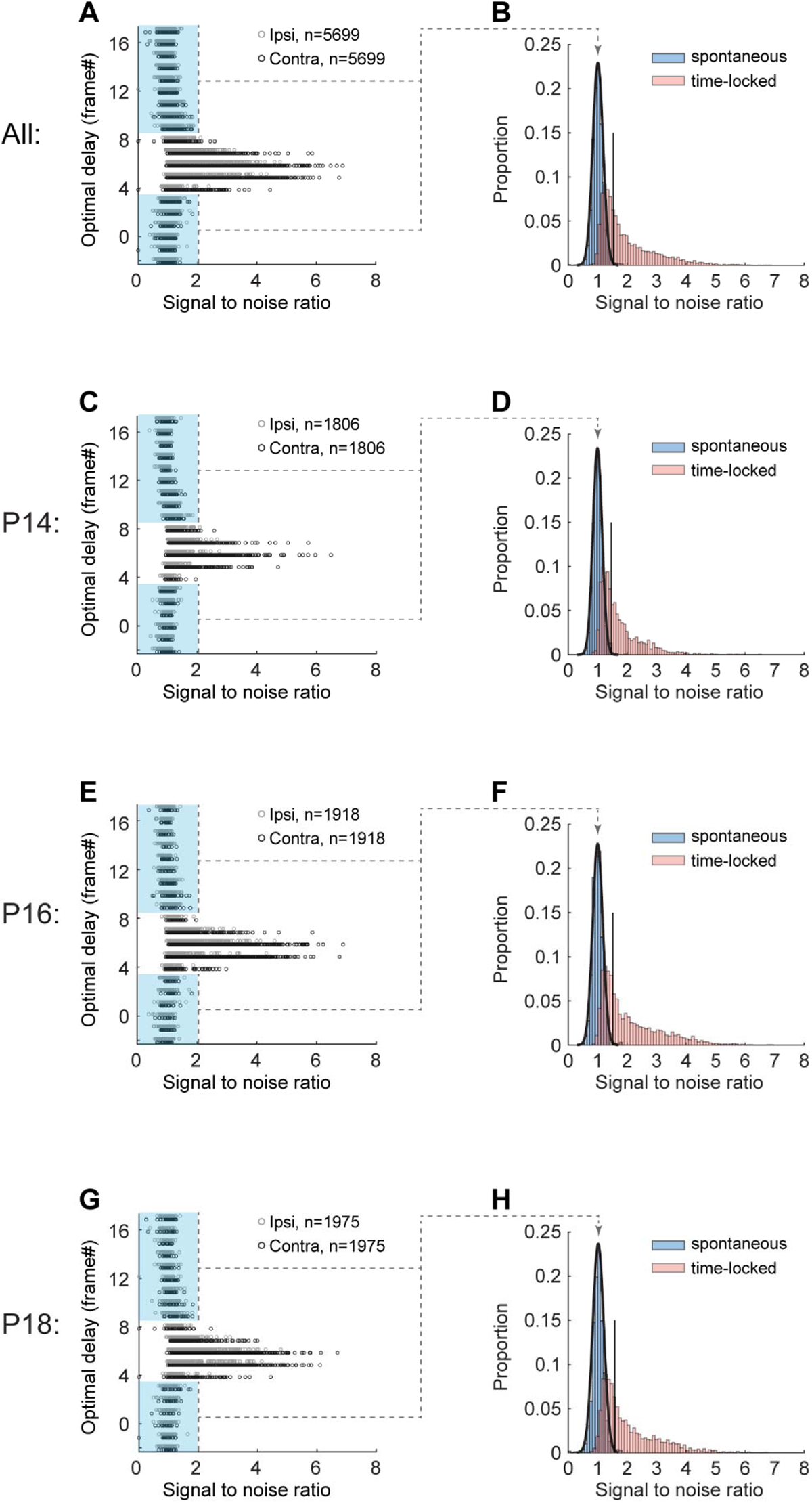
SNR distribution of neurons in three longitudinally imaged mice. Related to Figure 2. (A) Plot of SNR as a function of optimal delay for all 5699 layer 2/3 neurons imaged in P14, P16, and P18 mice in Figure 2. Dark and light gray represent responses to contralateral and ipsilateral eye stimulation, respectively. A cell with an optimal delay of -2 to 3 or 9 to 17 frames post stimulus onset was considered to be visually unresponsive (blue shading). (B) Blue shaded histogram and normal distribution fit of SNR values for unresponsive neurons. Black vertical line is 3 standard deviations above the mean of this normal distribution. Red shaded histogram: neurons whose spiking occurred between 4 and 8 frames post stimulus. Visually responsive neurons are those to the right of the vertical line, with optimal delays in this window that also had an SNR value above the threshold. (C–H) Same plots as in A and B but separated by age. The numbers of imaged neurons at each age are given above the plots in C, E, and G. Note that the distribution of SNR and the position of the black vertical line in B, D, F and H are highly similar.

**Figure S4.**
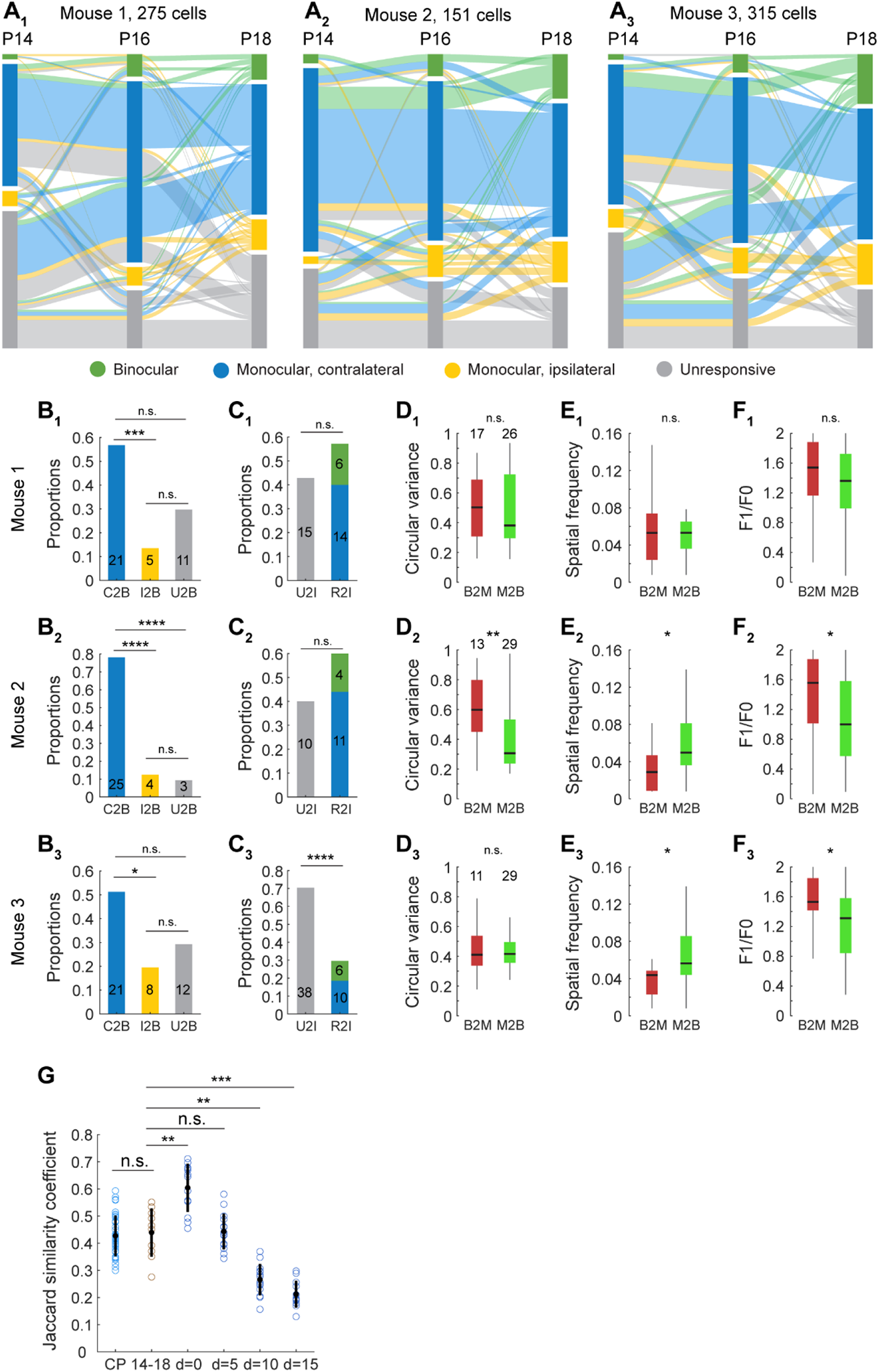
Longitudinal data sets from each mouse in Figure 2, and cell tracking quality of the longitudinal data. Related to Figure 2. (A1-A3) Sankey diagrams plotting the functional trajectory of neurons that were longitudinally imaged at P14, P16, and P18 in each mouse. The number of imaged neurons is given above each diagram. Cells are color coded as in Figure 2E. (B1-3-F1-3) As in Figure 2F-J. Sample numbers are shown in plots. Statistics: *, p<0.05; **, p<0.01; ***, p<0.001; ****, p<0.0001. (G) Cell tracking quality of the P14-16-18 longitudinal data is the same as the longitudinal tracking during the critical period. Plot of Jaccard similarity coefficients as a function of focal plane displacement (d=0 to d=15) (data from Tan et al., 2020); also plotted are these coefficients derived from the layer 2/3 neurons that were longitudinally imaged across the critical period (CP) (data from Tan et al., 2020), and these coefficients derived from the layer 2/3 neurons that were longitudinally imaged between P14 and 18 (14-18). Black dot and line indicate the mean and standard deviation. Number of data points from left to right are: 40, 10, 13, 15, 14 and 12. Mann-Whitney U test with Bonferroni correction was used for comparing between two groups of data. **, p<0.01; ***, p<0.001 after Bonferroni correction for pairwise comparison.

**Figure S5.**
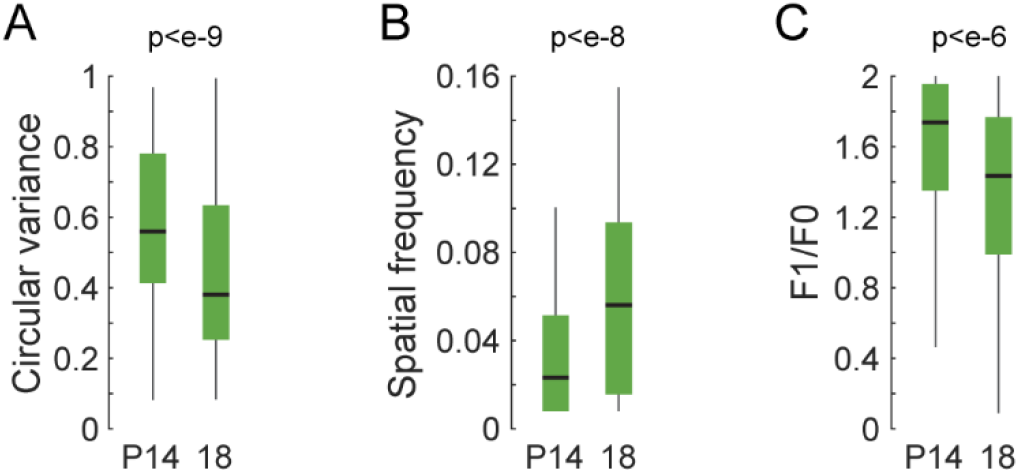
Acute measures of receptive field tuning of binocular neurons at eye opening and at P18. Related to Figure 2. (A-C) Measures of circular variance, spatial frequency preference, and complexity, respectively, made from 74 binocular neurons at P14 and 220 binocular neurons at P18 in normally reared mice. Note the improvement in tuning across all measures. Black horizontal line, median; box, quartiles with whiskers extending to 2.698σ. Mann-Whitney U test.

**Figure S6.**
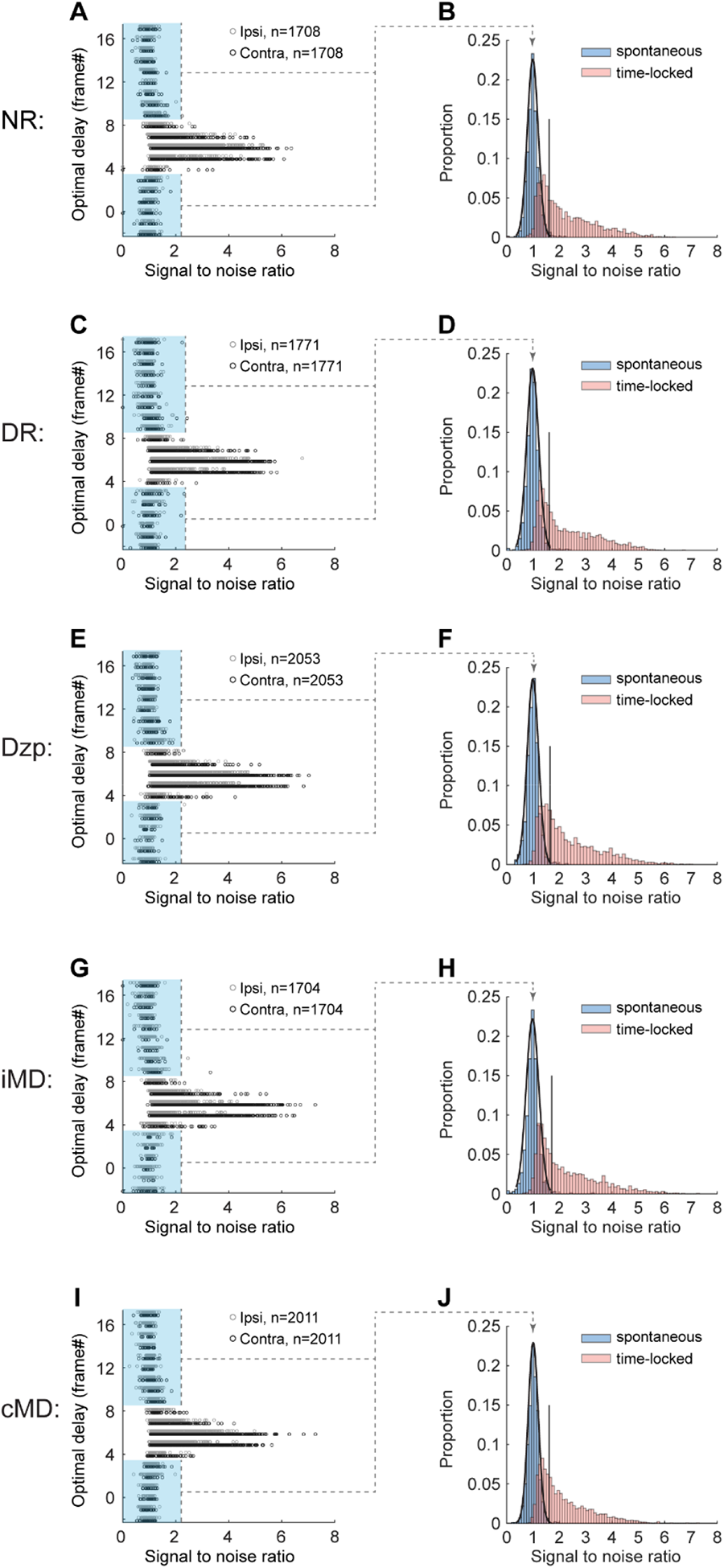
SNR distributions of all experience conditions at P18. Related to Figure 3-6. (A) Plot of SNR as a function of optimal delay for all 1708 L2/3 neurons imaged in four P18 normally reared (NR) mice. Dark and light gray represent responses to contralateral and ipsilateral eye stimulation, respectively. A cell with an optimal delay of -2 to 3 or 9 to 17 frames post stimulus onset was scored as visually unresponsive (blue shading). (B) Blue shaded histogram and normal distribution fit of SNR values for unresponsive neurons. Black vertical line is 3 standard deviations above the mean of this normal distribution. Red shaded histogram: neurons whose spiking occurred between 4 and 8 frames post stimulus. Visually responsive neurons are those to the right of the vertical line, with optimal delays in this window that also had an SNR value above the threshold. (C-J) Same plots as in A and B but separated by experience condition. The numbers of imaged neurons at each age are given above the plots in C, E, G and I. The numbers of mice are 4, 4, 4, and 3 for C, E, G and I, correspondingly. Note that the distribution of SNR and the position of the black vertical line in B, D, F, H and J are highly similar.

## STAR Methods

### RESOURCE AVAILABILITY

#### Lead Contact

Further information and requests for resources and reagents should be directed to and will be fulfilled by the Lead Contact, Joshua Trachtenberg.

#### Materials Availability

This study did not generate new unique reagents.

#### Data and Code Availability

Custom-written MATLAB scripts and data generated in this study are available from the Lead Contact upon request.

### EXPERIMENTAL MODEL AND SUBJECT DETAILS

All procedures were approved by UCLA’s Office of Animal Research Oversight (the Institutional Animal Care and Use Committee, IACUC) and were in accord with guidelines set by the US National Institutes of Health. Normally reared mice were housed in groups of 2-3 per cage in a reversed 12/12 light dark cycle. Dark-reared mice were housed in groups of 1-3 per cage in a light-tight cabinet that was additionally shielded with two layers of polyurethane-coated black nylon sheet (Thorlabs, BK5). Animals were naive subjects with no prior history of participation in research studies.

A total of 32 mice, both male (21) and female (11) were used in this study. In the studies of P14 mice involving flashed sinusoid gratings, 8 males and 5 females were used. P18 normally reared: 4 males. P18 dark reared from P12-P18: 3 males and 1 female. P18 with diazepam adminstered at P15 and 16: 3 males and 1 female (1 male and 1 female overlap with P14 flash grating). P18 with unilateral lid suture of the ipsilateral eye from P14-18: 2 males and 2 females (all overlap with P14 flash grating). P18 with unilateral lid suture to the contralateral eye from P14-18: 1 male and 2 females. P14 drifting grating: 1 male and 2 females. P14-18 longitudinally imaged: 2 males and 1 female.

Mice: All imaging was performed on mice expressing the slow variant of GCaMP6 in pyramidal neurons. These mice were derived from crosses of B6;DBA-Tg(tetO-GCaMP6s)2Niell/J (JAX Stock No: 024742)^52^ with B6;CBA-Tg(Camk2a-tTA)1Mmay/J (JAX Stock No: 003010)^53^. PCR was outsourced to Transnetyx (<u>transnetyx.com</u>).

### METHODS DETAILS

#### Surgery

All epifluorescence and two-photon imaging experiments were performed through chronically-implanted cranial windows as in Tan et al., 2020^6^. In brief, mice aged P10-P11 were administered with carprofen analgesia prior to surgery, anesthetized with isoflurane (5% for induction; 1.5–2% during surgery), mounted on a stereotaxic surgical stage via ear bars and a mouth bar and their body temperature maintained at 37°C via a heating pad. The scalp was removed, and the exposed skull was allowed to dry. The exposed skull and wound margins were then covered by a thin layer of Vetbond. Once dry, a thin layer of dental acrylic was applied around the wound margins, avoiding the region of skull overlying V1. A metal head bar was affixed with dental acrylic caudally to V1. A 3mm circular piece of skull overlying binocular V1 on the left hemisphere was removed using high-speed dental drill to first thin the bone along the circumference of this circle. Care was taken to ensure that the dura was not damaged at any time during drilling or removal of the skull. Once the skull was removed, a sterile 2.5 mm diameter cover glass was placed directly on the exposed dura and sealed to the surrounding skull with Vetbond. The remainder of the exposed skull and the margins of the coverglass were sealed with dental acrylic. Mice were then recovered on a heating pad. When alert, they were placed back in their home cage. Carprofen was administered daily for 3 days post-surgery. Mice were left to recover for at least 3 days prior to imaging.

#### Unilateral lid suture

The lid margins of one eye (ipsilateral or contralateral to the craniotomy, depending on the experiment) were sutured together at P14. Prior to doing so, the position of V1b was identified via retinotopic mapping of epifluorescent signals (see details below). In some mice, measures of receptive field tuning and binocularity were taken prior to lid suture via 2-photon imaging. For lid suture, mice were anesthetized with isoflurane (5% for induction; 2% during suturing). Two mattress sutures (5-0 vicryl) were placed through the lid margins on the left or the right eye.

After suture, the lid margins were inspected for any signs of gaps. Mice were recovered on heating pad before returning to their home cage. Thereafter, they were kept in normal rearing conditions and light/dark cycles. Sutures were checked daily to ensure the lid margins remained sealed. On the imaging day, sutures were removed, and the cornea was inspected for abrasions. Only mice whose sutured eye was open and clear after suture removal were used for 2-photon imaging. They were recovered for at least an hour in the dark after suture removal prior to imaging.

#### Diazepam injection

5 mg/ml diazepam (Hospira, Inc., NDC 0409-3213-12. Each mL contains 5 mg diazepam; 40% propylene glycol; 10% alcohol; 5% sodium benzoate and benzoic acid added as buffers and 1.5% benzyl alcohol added as a preservative. pH 6.6) was purchased, handled, and administered by UCLA DLAM. A day before injection at P14, retinotopic mapping was performed for each mouse to map V1b and confirm that neurons in visual cortex responded normally (see details below). In some mice, measures of receptive field tuning and binocularity were taken prior to diazepam injections via 2-photon imaging. Diazepam solution was administered intraperitoneally at a dose of 30 mg/kg^36, 37^ at P15 and again at P16. Mice fell into a stupor one minute after injection and this lasted for several hours. Diazepam injected mice lost 4-9% of body weight over the three days after the first injection. To keep mice hydrated and to maintain body weight, we administered approximately 7% body weight of lactated Ringer’s solution subcutaneously from the first day of diazepam injection to two days after the second diazepam dose.

#### Mapping of binocular area of the primary visual cortex

The binocular area of primary visual cortex on the left hemisphere for each mouse was identified using low magnification, epifluorescence imaging of GCaMP6s signals as in Tan et al., 2020^6^. For all experiments except P18 NR and DR, visual areas were mapped at P14. For P18NR mice, visual areas were mapped at P17. For P18 DR mice, visual areas were mapped about an hour prior to 2P imaging. Briefly, GCaMP6s was excited using a 470nm light-emitting diode. A 27-inch LCD monitor (ASUS) was positioned such that the binocular visual field fell in the center of the monitor. The screen size was 112 deg in azimuth and 63 deg in elevation and the monitor was placed 20 cm from the eyes. We presented a contrast reversing checkerboard (checker size 10×10 degree) windowed by a 1D Gaussian along the horizontal or vertical axis to both eyes. The checkerboard drifted normal to its orientation and swept the full screen width in 10 sec. We used both directions of motion to obtain an absolute phase map along the two axes.

Eight cycles were recorded for each of the four cardinal directions. Images of were acquired at 10 frames per second with a PCO edge 4.2 sCMOS camera. We used a 35mm fixed focal length lens (Edmund optics, 35mm/F1.65, #85362, 3mm field of view) in place of the standard objective lens on our microscope for macroscopic imaging. The camera focused on the pia surface. Retinotopic maps of azimuth and elevation were generated as in Kalatsky and Stryker, 2003^54^, and visual field sign maps were calculated as in Garrett et al., 2014^55^. The binocular area of the primary cortex was defined as the region of primary visual cortex adjacent to the higher visual area LM (**Figure 1B**).

#### Two photon calcium imaging

2-photon imaging was performed as in Tan et al., 2020^6^. Briefly, imaging was targeted to the binocular area of V1 using a resonant/galvo scanning two-photon microscope (Neurolabware, Los Angeles, CA) controlled by Scanbox image acquisition software (Los Angeles, CA). A Coherent Discovery TPC laser (Santa Clara, CA) running at 920 nm focused through a 16x water-immersion objective lens (Nikon, 0.8 numerical aperture) was used to excite GCaMP6s. The objective was set at an angle of ∼10 degrees from the plumb line to reduce the slope of the imaging planes. Image sequences (512×796 pixels, 490×630 μ were captured at 15.5 Hz at a depth of 120 to 300μ below the pial surface on alert, head-fixed mice that were free to run on a 3D-printed running wheel (14cm diameter). A rotary encoder was used to record the rotations of this running wheel. Eye movements and changes in pupil size were recorded using a Dalsa Genie M1280 camera (Teledyne Dalsa) fitted with a 740nm long-pass filter. The eye tracking camera was triggered at scanning frame rate. To measure responses of neurons to each eye separately, an opaque patch was placed immediately in front of one eye when recording neuronal responses to visual stimuli presented to the other eye.

#### Visual stimulation during 2-photon imaging

Based on retinotopic mapping results, the position of the 27-inch LCD monitor was adjusted slightly so that the receptive fields of the imaging field of view were located on the center of the screen. The monitor was placed 20 cm from the eyes and covered 112 degrees in azimuth and 63 degrees in elevation. Screen refreshed at 60 Hz and stimuli were shown in full screen with contrast set at 80%.

##### Flash sinusoidal gratings

Flash gratings were the same as in Tan et al., 2020^6^. Briefly, a set of sinusoidal gratings with 18 orientations (equal intervals of 10 degrees from 0 to 170 degrees), 12 spatial frequencies (equal steps on a logarithmic scale from 0.0079 to 0.1549 cycles per degree) and 8 spatial phases were generated in real-time by a Processing sketch using OpenGL shaders (see https://processing.org). These static gratings were presented at 4 Hz in pseudo-random sequence. Imaging sessions were 15 min long (3600 stimuli in total), thus each combination of orientation and spatial frequency appeared 16 or 17 times. Each of the 8 spatial phases for an orientation/spatial frequency combination appeared twice (F1/F0 values were calculated using responses of neurons as a function of spatial phase). Transistor-transistor logic signals were used to synchronize visual stimulation and imaging data. The stimulus computer generated these signals and these were sampled by the microscope electronics and time-stamped by the acquisition computer to indicate the frame and line number being scanned at the time of the TTL.

##### Drifting sinusoidal gratings

A set of sinusoidal gratings with 16 directions (equal intervals of 22.5 degrees from 0 to 337.5 degrees) and 2 spatial frequencies (0.03 and 0.13 cycles per degree, for low and high spatial frequencies) were generated in real-time by a Processing sketch using OpenGL shaders (see https://processing.org). The full set of gratings (32 combinations of directions and spatial frequencies) were shown 5 times in one experiment. Stimuli were presented in pseudo-random sequence, with each grating drifting at 1 Hz for 2 seconds, followed by 4 seconds of gray screen. Thus, the total time for one experiment is 16 min. Transistor-transistor logic signals were generated by the stimulus computer at the onset and closure of each drifting stimulus. These signals were sampled by the microscope and time-stamped with the frame and line number being scanned at that time to synchronize visual stimulation and imaging data.

#### Analysis of two-photon imaging data

##### Image processing

The pipeline for image processing is described in detail in Tan et al., 2020^6^. Briefly, movies from the same plane for each eye were concatenated and motion corrected. Regions of interest (ROI) corresponding to pyramidal neuron soma were determined using a Matlab graphical user interface tool (Scanbox, Los Angeles, CA). Using this GUI, we computed pixel-wise correlations of fluorescence changes over time with 1200 evenly spaced imaging frames (4% of all frames). The temporal correlation of pixels was used to determine the boundary of ROI for each neuron. Data from all experiments except the P14-18 longitudinal imaging were aligned and segmented in this way. Data from the longitudinal experiment were aligned and segmented using suite2p^56^. ROIs determined for each experiment were inspected and confirmed visually via superposition with mean fluorescence image. Control analysis was carried out to confirm that ROIs determined by both Scanbox pipeline and suite2p for the same experiment rendered the same outcomes. After segmentation, the fluorescence signal for each ROI and surrounding neuropil was extracted.

The neuropil signal for a given ROI was computed by dilating the ROI with a disk of 8-pixel radius. The original ROI and those of other cells that overlap with the region were excluded, and the average signal within this area was computed. The signal obtained from the ROI was then robustly regressed on the neuropil. The residual represents the corrected signal of the ROI. The correction factor results from the slope of the robust regression. Neuronal spiking was estimated via non-negative temporal deconvolution of the corrected ROI signal using Vanilla algorithm^57^. Subsequently, fluorescent signals and estimated spiking for each cell were split into separate files corresponding to the individual imaging session for each eye. Each imaging experiment was independently segmented.

##### Calculation of response properties in flash grating experiments

The analysis for flash grating experiments, including SNR calculation, tuning kernels of orientation and spatial frequency responses, phase invariance, orientation and spatial frequency preference, binocular matching coefficients were described in detailed in Tan et al., 2020^6^. They are described here in brief.

###### Identification of visually responsive neurons using SNR

Signal to noise ratio (SNR) was used to identify neurons with significant visual responses. SNR for each neuron was calculated based on the optimal delay (the imaging frame after stimulus onset at which neuron’s GCaMP6s fluorescence reaches maximum), with signal being the mean of spiking standard deviation at the optimal delay (5-7 frames, thus ∼0.387 sec, after stimulus onset), and noise as this value at frames well before or after stimulus onset (frames –2 to 0, and 13 to 17). Neurons whose optimal delays occurred outside of the time-locked stimulus response window of 4 to 8 frames (**Figure S1, S3, S6**, blue highlight) were spontaneously active but visually unresponsive. They had SNR values close to 1. The SNR values of these unresponsive neurons were normally distributed (mean = 1.0) over a narrow range (**Figure S1, S3, S6**, blue shaded histogram). Spontaneously active neurons with optimal delays naturally occurring in the 4-8 frame time window (**Figure S1, S3, S6**, red shaded) can be distinguished from visually responsive neurons by SNR. This SNR threshold is defined at 3 standard deviations above the mean SNR of the blue shaded normal distribution (**Figure S1, S3, S6**, vertical black line). Visually responsive neurons had optimal delays between frames 4 and 8, and SNRs greater than this threshold (**Figure S1C**). SNR values were calculated separately for responses to the ipsilateral or contralateral eye.

###### Tuning kernel for orientation and spatial frequency

The estimation of the tuning kernel was performed by fitting a linear model between the response and the stimulus^58^. Cross-correlation maps were used to show each neuron’s spiking level to each visual stimulus (orientation and spatial frequency), and were computed by averaging responses over spatial phases. The final tuning kernel of a neuron was defined as the correlation map at the optimal delay (**Figure 1D**).

###### F1/F0 measurement for phase invariance

F1/F0 is the ratio of the 1^st^ Fourier harmonic and 0^th^ Fourier harmonic for a given cell across different spatial phases ^59^. For complex cells the F1/F0<1, while for simple cells F1/F0>1^60^.

###### Orientation and spatial frequency preference

We used horizontal (for spatial frequency) and vertical (for orientation) slices of the tuning kernel through the peak response to calculate orientation and spatial frequency preferences.

Orientation preference calculation:

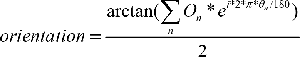

O_n_ is a 1×18 array, in which a level of estimated spiking (O_1_ to O_18_) occurs at orientations θ_n_ (0 to 170 degrees, spaced every 10 degrees). Orientation is calculated in radians and then converted to degrees.

Spatial frequency preference calculation:

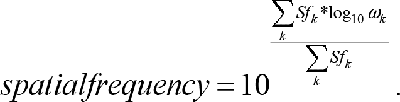

Sf_k_ is a 1×12 array, in which a level of estimated spiking (Sf_1_ to Sf_12_) occurs at spatial frequencies ω_k_ (12 equal steps on a logarithmic scale from 0.0079 to 0.1549 cycles per degree).

###### Circular variance

Circular variance is a measure of orientation selectivity, with limits from zero to one. The circular variance of a neuron whose estimated spiking, O_n_, occurred at orientations θ_n_ (0° to 170°, spaced every 10°), is defined by

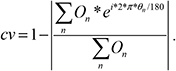

Neurons with higher orientation selectivity have lower circular variance values (see examples in Figure 1E).

###### Binocular matching coefficient

This was defined as the correlation coefficient between contralateral and ipsilateral tuning kernels of binocular neurons.

##### Calculation of response properties in drifting grating experiments

In each experiment, the GCaMP6s fluorescence signal of each cell was derived by subtracting the neuropil signal by the ROI signal. This cellular fluorescence, F, is then split into signal segments corresponding to all stimuli. Each segment includes signals from two seconds before stimulus onset to four seconds after stimulus closure for its corresponding drifting grating stimulus. The baseline F_0_ for each segment was defined by the mean signal of 16 imaging frames before stimulus onset. The baseline corrected activity for each segment, dF/F_0_, was calculated as (F-F_0_)/F_0_. Stimulus-evoked responses was then defined as the dF/F_0_ from 6 imaging frames after the stimulus onset to 6 frames after stimulus closure. Noise was defined by the dF/F_0_ in the 4^th^ second after stimulus closure and was pooled across all stimuli in an experiment. Trial-averaged responses were calculated by taking the mean of stimulus-evoked responses across repeated trials to the same stimulus. The above-mentioned process was done separately for recordings to the contralateral or ipsilateral eye. Cells were considered monocular if the trial-averaged response to any stimulus direction was greater than two standard deviations above the noise mean for the contralateral or the ipsilateral eye^61^. Cells were considered binocular if the trial-averaged response to any stimulus direction was greater than two standard deviations above the noise mean for each eye.

For responsive cells, the preferred spatial frequency (either 0.03 or 0.13 cycles per degree) is at which cells have highest trial averaged response to a direction.

Orientation preference calculation:

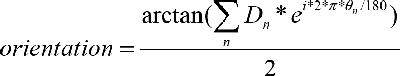

Circular variance calculation:

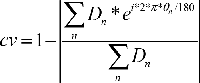

D_n_ is a 1×16 array, in which trial-averaged response (D_1_ to D_16_) occurs at directions θ_n_ (0 to 337.5 degrees, spaced every 22.5 degrees). Orientation was calculated in radians and subsequently converted to degrees.

##### Longitudinal imaging and analysis

Longitudinal imaging and analysis were performed as in Tan et al., 2020^6^. Briefly, the same objective angle was used to locate the same imaging plane across days. Images acquired on the previous imaging day are used as reference. Fine adjustment of imaging depth was made by manual z-scanning in small steps of 2 μ for ∼20 μ while running visual stimulation to identify the same imaging plane that matched the reference. Imaging planes were considered to be near identical when cell morphology across the four quadrants of the image matched those in the reference. To identify neurons tracked between adjacent days, a control point-based affine geometric transformation (Matlab syntax: cpselect, fitgeotrans and imwarp) was performed to correct the plane rotation. This method can work for both plane rotation and size differences. In our experiment, we only witnessed plane rotation. There was no obvious growth along the x-, y- or z-axis. The overlap of ROIs between adjacent imaging days was calculated. If the overlapping area between two ROIs was bigger than 50% of their union areas, the two ROIs were considered to be the same cell. The quality of longitudinal imaging in this study is identical to that in Tan et al., 2020^6^ (**Figure S4G**).

##### Figure plotting

Sankey diagrams in Figure 2E and Figure S4A were plotted using RStudio with dplyr and alluvial libraries.

#### Quantification and Statistical Analysis

A power analysis was not performed a-priori to determine sample size. All statistical analyses were performed in MATLAB (https://www.mathworks.com/), using non-parametric tests with significance levels set at α < 0.05. Bonferroni corrections for multiple comparisons were applied when necessary. Mann-Whitney U-tests (Wilcoxon rank sum test) were used to test differences between two independent populations. When comparing more than two populations that were non-normally distributed, a Kruskal-Wallis test, a nonparametric version of one-way ANOVA, was used. Where significant differences were found, post hoc Mann-Whitney U-tests with Bonferroni correction were used to test for significant differences between within group pairs. For pairwise comparisons of proportions, Chi-square test was used (https://www.mathworks.com/matlabcentral/fileexchange/45966-compare-two-proportions-chi-square).

